# Bradykinin receptor expression and bradykinin-mediated sensitization of human sensory neurons

**DOI:** 10.1101/2023.03.31.534820

**Authors:** Jiwon Yi, Zachariah Bertels, John Smith Del Rosario, Allie J. Widman, Richard A. Slivicki, Maria Payne, Henry M. Susser, Bryan A. Copits, Robert W. Gereau

**Affiliations:** Department of Anesthesiology, Washington University Pain Center, Washington University School of Medicine, St. Louis, MO, United States; Neuroscience Graduate Program, Division of Biology & Biomedical Sciences, Washington University School of Medicine, St. Louis, MO, United States; Department of Neuroscience, Washington University, St. Louis, MO, United States; Department of Biomedical Engineering, Washington University, St. Louis, MO, United States

**Keywords:** bradykinin, human dorsal root ganglia, sensory neurons, pain

## Abstract

Bradykinin is a peptide implicated in inflammatory pain in both humans and rodents. In rodent sensory neurons, activation of B1 and B2 bradykinin receptors induces neuronal hyperexcitability. Recent evidence suggests that human and rodent dorsal root ganglia (DRG), which contain the cell bodies of sensory neurons, differ in the expression and function of key GPCRs and ion channels; whether BK receptor expression and function are conserved across species has not been studied in depth. In this study, we used human DRG tissue from organ donors to provide a detailed characterization of bradykinin receptor expression and bradykinin-induced changes in the excitability of human sensory neurons. We found that B2 and, to a lesser extent, B1 receptors are expressed by human DRG neurons and satellite glial cells. B2 receptors were enriched in the nociceptor subpopulation. Using patch-clamp electrophysiology, we found that acute bradykinin increases the excitability of human sensory neurons, while prolonged exposure to bradykinin decreases neuronal excitability in a subpopulation of human DRG neurons. Finally, our analyses suggest that donor’s history of chronic pain and age may be predictors of higher B1 receptor expression in human DRG neurons. Together, these results indicate that acute BK-induced hyperexcitability, first identified in rodents, is conserved in humans and provide further evidence supporting BK signaling as a potential therapeutic target for treating pain in humans.

## Introduction

Bradykinin (BK) is a potent inflammatory mediator released during tissue injury and inflammation [4,17,19,32]. BK is associated with pathogenesis of inflammatory pain in both humans and rodents: intradermal injection of BK in healthy human volunteers or intraplantar injection of BK in rodents can lead to local inflammation and cutaneous thermal hyperalgesia [33,49]. Systemic and local administration of B1 and/or B2 receptor antagonists can be anti-nociceptive in neuropathic, visceral, inflammatory, and other rodent pain models [10,16,18,38,40,59]. Additionally, patients with painful disorders like arthritis, endometriosis, and myalgia exhibit elevated levels of BK in damaged tissue and/or plasma compared to healthy controls, with level of kinin correlating with pain degree and/or disease progression [4,6,17,21,29,37,41].

BK-mediated hyperalgesia is thought to occur, in part, through the activation of Gq-coupled B1 and B2 receptors expressed on sensory neurons. In rodent dorsal root ganglion (DRG) neurons *in vitro*, acute application of BK can induce inward current, action potential firing, hyperexcitability, and TRPV1 sensitization [5,8,9,12,28,34–36,54]. While B2 is the predominant BK receptor subtype in the DRG of uninjured animals [12], B1 receptors can be synthesized *de novo* after formalin injection, nerve injury, or exposure to inflammatory growth factors, thereby further contributing to BK-mediated pain [12,26,38,57,59]. Additionally, satellite glial cells in the DRG and trigeminal ganglia express BK receptors and may contribute to BK’s effects on neuronal physiology in the periphery [7,20]. Together, these preclinical studies indicate that BK receptor activation on sensory neurons is integral to development and/or maintenance of pain.

While the mechanism underlying BK-induced pain and hyperalgesia has been extensively studied in rodent models, it is yet unclear whether the same mechanism translates across species to humans. Of note, an increasing number of studies has demonstrated that expression and function of key ion channels and receptors differ between human and rodent DRG [11,14,39,46,48,50,51,55]. This highlights the need for validating targets in human tissue to maximize the translational potential of preclinically-identified analgesics. Whole-tissue transcriptomics shows that the B2 receptor is expressed in human DRG (hDRG) [46], and acute application of BK elicits spontaneous activity and induces hyperexcitability in a subset of human DRG neurons [11]. However, whether BK receptors are expressed among human nociceptor subpopulations, and whether BK modulates all physiological subtypes of hDRG [47] is yet unknown.

In the present study, we evaluated the expression of B1 (*BDKRB1*) and B2 (*BDKRB2*) receptors in hDRG tissue obtained postmortem from organ donors. We additionally examined the effects of acute and prolonged exposure to BK on the excitability of two physiologically distinct subpopulations of human sensory neurons. Finally, we assessed whether donors’ pain history is associated with changes in BK receptor expression or functional modulation of neuronal excitability. We found that human DRG neurons express functional BK receptors, whose modulation of neuronal excitability is dependent on both the duration of BK exposure and physiological subtype of the sensory neuron. These findings suggest that BK-induced acute hyperexcitability is conserved from rodents to humans and provide further evidence of BK signaling as a viable analgesic target.

## Results

### Bradykinin receptors are expressed in human sensory neurons

Previously, a whole-tissue transcriptomics study of hDRG has found that B2 and, to a lesser extent, B1 receptors are expressed in the hDRG [46]. To characterize the distribution pattern of BK receptors in hDRG subpopulations, we performed RNAscope *in situ* hybridization for *BDKRB1* and *BDKRB2*, the genes for B1 and B2 receptors, respectively, on DRG samples from donors without pain history (Fig 1A, B; Table 2). Consistent with the findings in whole-tissue transcriptomics study of human and mouse DRG[46], *BDKRB1* was detected at low levels and in a small fraction (<10%) of hDRG neurons (Fig 1C, D), while *BDKRB2* was detected in a larger fraction (48-61%; Fig 1C) of hDRG neurons and at significantly higher levels compared to *BDKRB1* (Fig 1D; η^2^=0.268, d=1.211). More than half of *BDKRB1*-expressing neurons were medium-to large-sized neurons with diameter greater than 40µm (Fig 1D), while *BDKRB2* was expressed across sensory neurons of varying sizes (16um-131um) (Fig 1F). The DRG neuron size distribution was consistent between donors (data not shown).

**Table 1.**
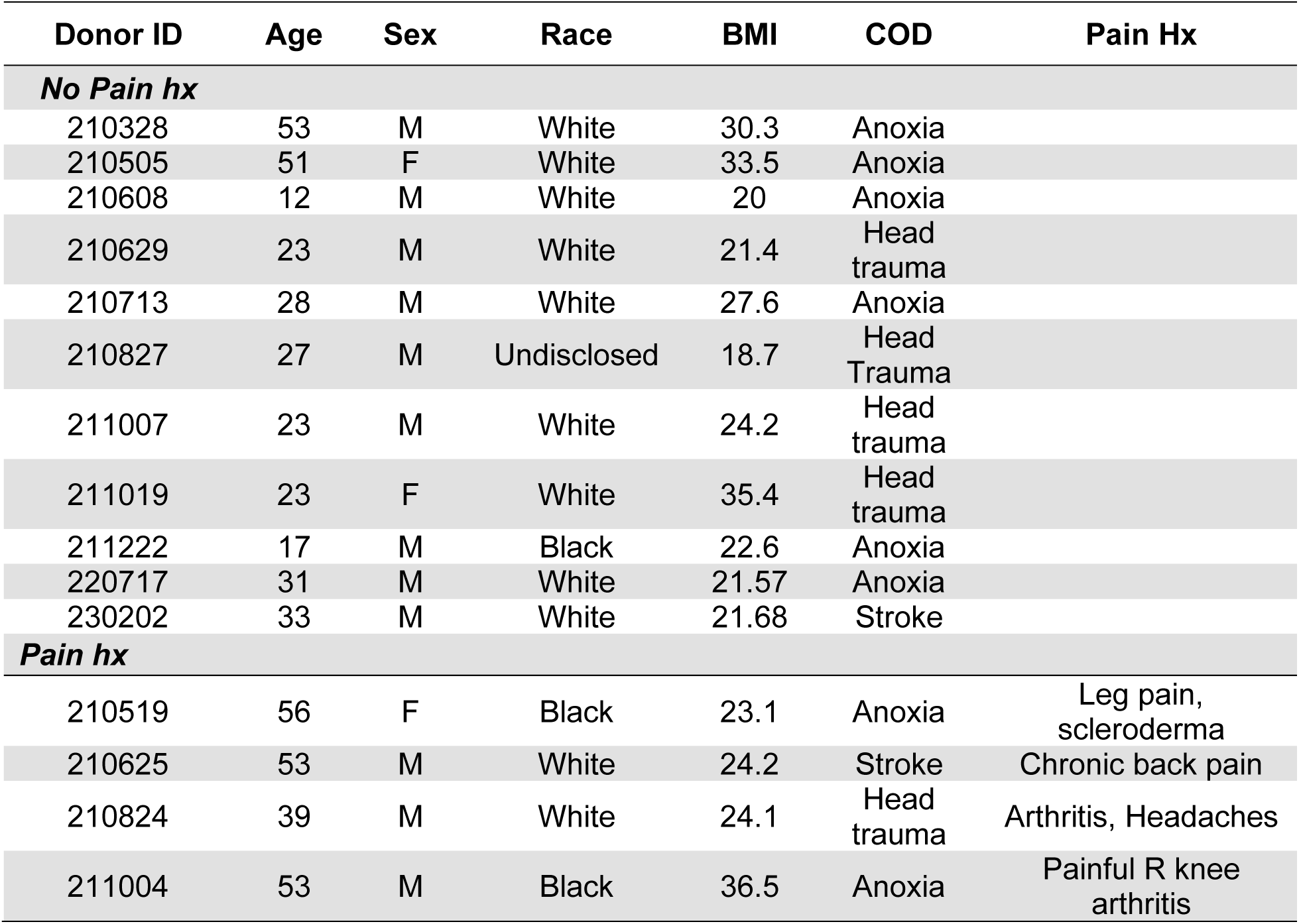
Demographic information and medical history of donors included in physiology experiments.

| Donor ID | Age | Sex | Race | BMI | COD | Pain Hx |
| --- | --- | --- | --- | --- | --- | --- |
| ***No Pain hx*** |  |  |  |  |  |  |
| 210328 | 53 | M | White | 30.3 | Anoxia |  |
| 210505 | 51 | F | White | 33.5 | Anoxia |  |
| 210608 | 12 | M | White | 20 | Anoxia |  |
| 210629 | 23 | M | White | 21.4 | Head trauma |  |
| 210713 | 28 | M | White | 27.6 | Anoxia |  |
| 210827 | 27 | M | Undisclosed | 18.7 | Head Trauma |  |
| 211007 | 23 | M | White | 24.2 | Head trauma |  |
| 211019 | 23 | F | White | 35.4 | Head trauma |  |
| 211222 | 17 | M | Black | 22.6 | Anoxia |  |
| 220717 | 31 | M | White | 21.57 | Anoxia |  |
| 230202 | 33 | M | White | 21.68 | Stroke |  |
| ***Pain hx*** |  |  |  |  |  |  |
| 210519 | 56 | F | Black | 23.1 | Anoxia | Leg pain, scleroderma |
| 210625 | 53 | M | White | 24.2 | Stroke | Chronic back pain |
| 210824 | 39 | M | White | 24.1 | Head trauma | Arthritis, Headaches |
| 211004 | 53 | M | Black | 36.5 | Anoxia | Painful R knee arthritis |

**Table 2.** Demographic information and medical history of donors included in RNAScope experiments.

| Donor ID | Age | Sex | Race | COD | Level | Pain Hx |
| --- | --- | --- | --- | --- | --- | --- |
| ***No Pain hx*** |  |  |  |  |  |  |
| 20210329 | 53 | M | White | Anoxia | L3 |  |
| 20210706 | 45 | M | White | Anoxia | L1 |  |
| 20211222 | 17 | M | Black | Anoxia | L4 |  |
| ***Pain hx*** |  |  |  |  |  |  |
| 20210404 | 23 | M | White | Head trauma | L1 | Chronic joint pain |
| 20210813 | 32 | M | White | Head trauma | L2 | Frequent joint pain in hands |
| 20220916 | 58 | F | Unknown | Stroke | Lumbar (unspecified) | Painful degenerative disc disease |

**Figure 1.**
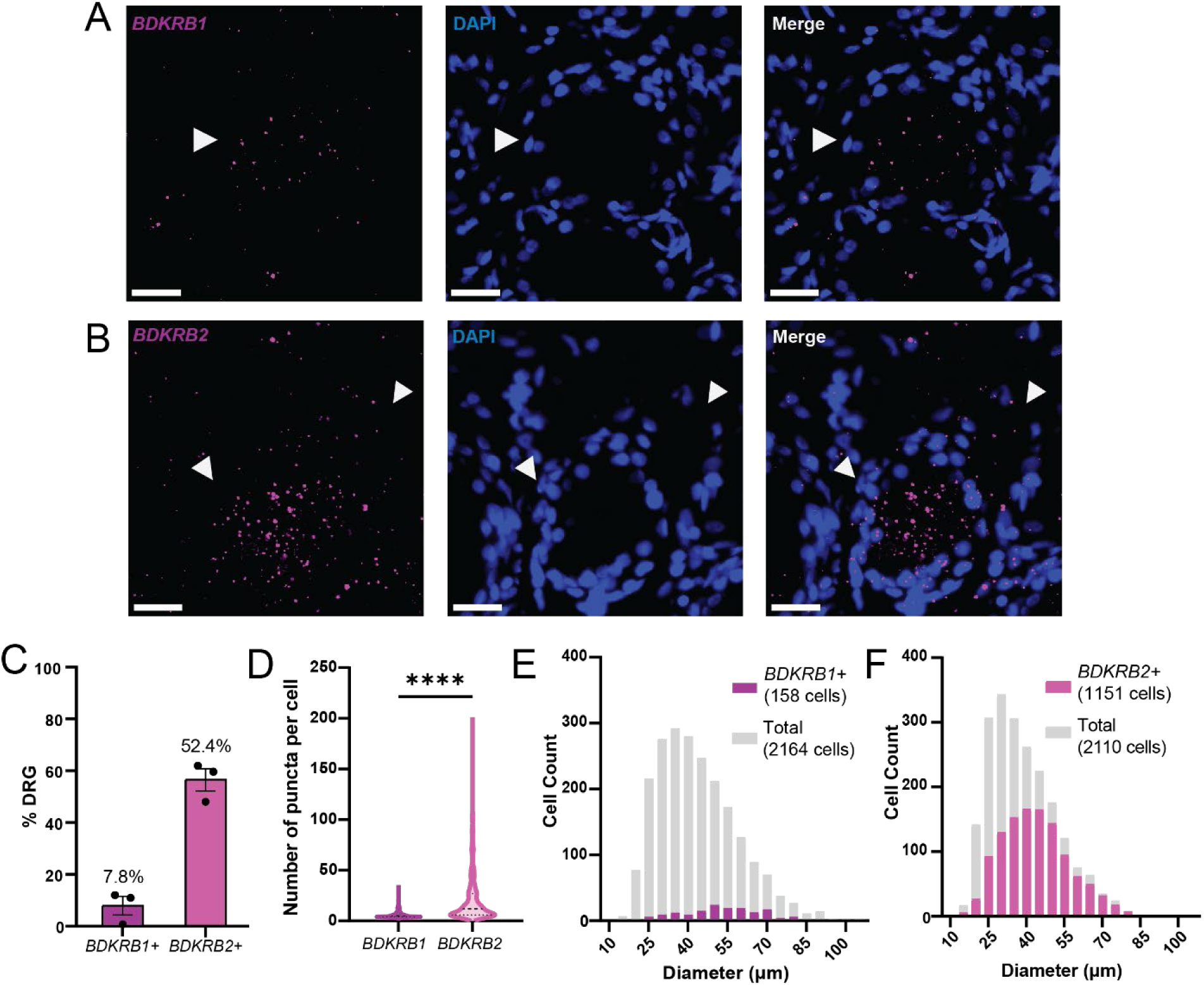
B1 and B2 bradykinin receptors are expressed by human sensory neurons. A) Representative image of *BDKRB1* expression in human DRG assessed using RNAscope; scale bar, 25µm. Arrowheads mark *BDKRB1*-expressing hDRG neuron. B) Representative image of *BDKRB2* expression in human DRG assessed using RNAscope; scale bar, 25m. Arrowheads mark *BDKRB2*-expressing neurons. C) *BDKRB1* and *BDKRB2* expression in human DRG averaged across donors (N=3 donors, 2-4 sections, 323-1044 cells per donor). D) Count of RNA puncta per cell in *BDKRB1*+ (left; n=405) or *BDKRB2*+ (right; n=655) hDRG neurons; ****p<0.0001, Mann-Whitney test. E) Size distribution of *BDKRB1*-expressing neurons (purple) compared to all DRG neurons (grey; n=2164 total neurons, 158 *BDKRB1*+ neurons from 3 donors). F) Size distribution of *BDKRB2*-expressing neurons (magenta) compared to all DRG neurons (grey; n=2110 total neurons, 1151 *BDKRB1*+ neurons from 3 donors). Data represent mean ± s.e.m.

To assess whether *BDKRB2* is specifically expressed by nociceptors in hDRG, we examined whether *BDKRB2* mRNA is localized in cells expressing *SCN10A*. *SCN10A* was chosen as a marker for human nociceptors based on a previously published spatial transcriptomics study, which demonstrated that *SCN10A* is specifically and highly enriched among multiple nociceptor subpopulations in hDRG[55]. *BDKRB2* and *SCN10A* were co-expressed in 1/3 of all sensory neurons analyzed (31.3%, n=2663 cells; Fig 2A, B). A majority (72.4%) of *BDKRB2*-expressing neurons also expressed *SCN10A*, indicating that *BDKRB2*-expressing cells are predominantly nociceptors (Fig 2C). Among the *SCN10A*-expressing putative nociceptors, around half (51.6%) also expressed *BDKRB2* (Fig 2C). Interestingly, we also observed that *BDKRB2* co-localized with FABP7, a marker of satellite glial cells (Fig 2D). We were unable to quantify the relative proportion of BDKRB2-expressing satellite glial cells, as segmenting individual satellite glial cells that have been imaged using fluorescence or confocal microscopy is challenging due to the tight contact between the multiple satellite glial cells that surround a hDRG neuron [1]. Together, these results suggest that B2 receptor is expressed in both nociceptor and glial subpopulations in the DRG.

**Figure 2.**
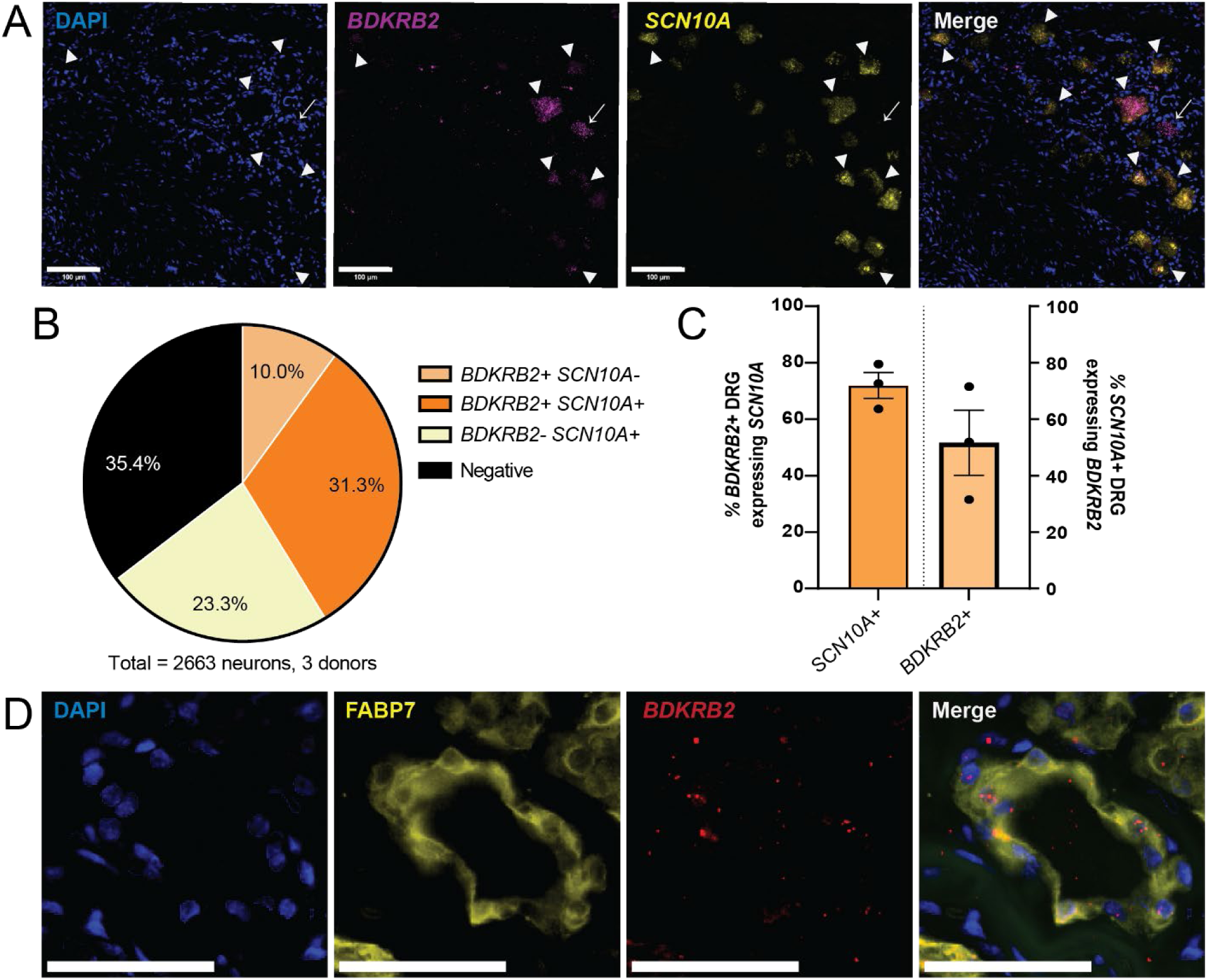
B2 bradykinin receptor is expressed in human nociceptors and satellite glial cells. A) Representative image of human DRG expressing mRNA for *BDKRB2* (magenta) and *SCN10A* (yellow), the human nociceptor marker. Arrowheads mark cells co-expressing *BDKRB2* and *SCN10A*; arrows mark cell that expresses *BDKRB2*, but not *SCN10A*; scale bar, 100 µm. B) Distribution of *BDKRB2* and *SCN10A* across human sensory neurons analyzed (2663 neurons from 3 donors). C) Quantification of BDKRB2 distribution, averaged across donors (N=3 donors, 2-3 sections and 338-1349 neurons per donor). Left, proportion of *BDKRB2*-expressing neurons that co-express SCN10A. Right, proportion of *SCN10A*-expressing nociceptors that express *BDKRB2*. D) Representative image of *BDKRB2* expression in FABP7-expressing satellite glial cells in human DRG; scale bar, 50 µm. Data represent mean ± s.e.m.

### Acute bradykinin sensitizes human DRG neurons

In a previous study, we reported that acute application of BK can induce spontaneous activity and reduce rheobase in hDRG neurons [11]. Additionally, recent work has suggested that both rodent and human sensory neurons exist in distinct physiological phenotypes that can be distinguished by single- or repetitively firing discharge pattern in response to a stepwise current injection [3,43,47]. Therefore, we tested whether acute BK-induced hyperexcitability is consistent across small-diameter neurons with different physiological phenotypes.

We found that acute (2-3 min) application of 100nM BK led to spontaneous activity or depolarization in 75% of neurons recorded (Fig 3A, B; 37.01 ± 7.829 µm, mean diameter ± SD). The depolarization in resting membrane potential was seen in both single- and repetitive-spiking hDRG (Fig 3C). Additionally, 72% of all neurons displayed either reduced rheobase or increased spike firing (Fig 3D, E), which contributed to a leftward-shift in the input-output curve of number of spikes fired in response to increasing current steps in both pooled neurons and the single-spiker subgroup (Fig 3F). Acute BK additionally led to increased input resistance and reduced rheobase in single spikers, consistent with hyperexcitability (Table 3). While BK treatment led to increased spike firing in many repetitively firing neurons, statistical analysis showed no significant effect of treatment, likely due to the variability in spike frequency between repetitive-firing neurons. Together, these results indicate that acute BK application can increase the intrinsic excitability of human sensory neurons.

**Figure 3.**
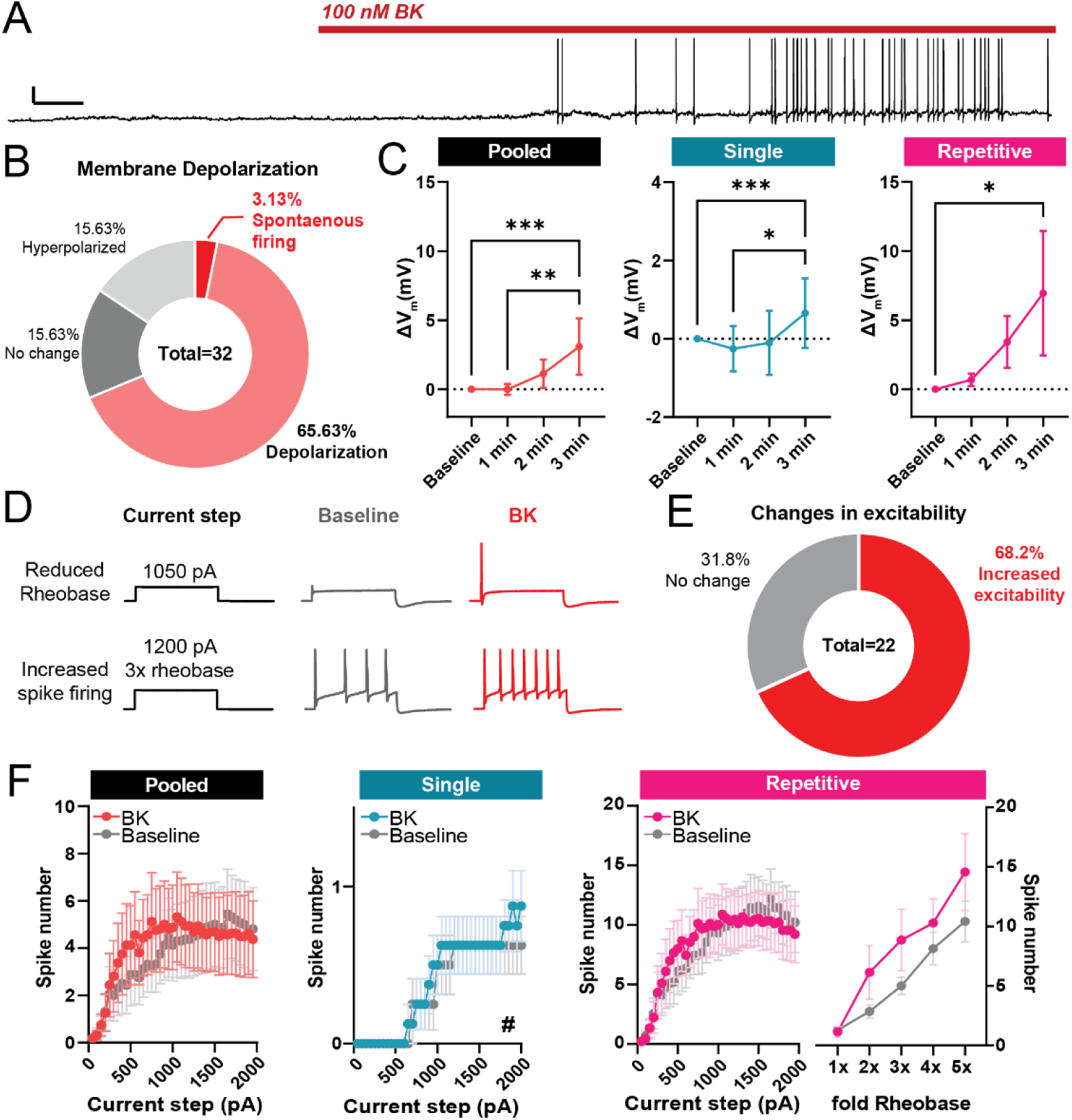
Acute exposure to BK sensitizes small-diameter human DRG neurons. A) Representative voltage trace of a sensory neuron with spontaneous action potential firing during bath application of BK (100 nM); Scale bars, 20mV/10s. B) Distribution of neuronal responses to acute BK based on resting membrane potential depolarization and spontaneous activity during the first 2 minutes of application. C) Changes in resting membrane potential in human DRG neurons grouped by firing phenotype. Left, pooled analysis of all recorded neurons (n=19 cells); middle, analysis of single-spiking neurons only (n=11); right, analysis of repetitively-firing neurons only (n=8). *p<0.05 **p<0.01, ***p<0.001, Friedman’s test with Dunn’s post-hoc multiple comparison test. D) Representative traces reduction in rheobase (top) and increase in number of spikes fired during suprathreshold current injections (bottom) following treatment with 100nM BK. E) Distribution of neuronal response to acute BK based on shifts in excitability, as defined by a leftward shift in the input-output curve. F) Input-output curve of BK-sensitive human DRG neurons, before and during acute bath application of BK. Left, pooled analysis of all recorded neurons (n=16 cells); middle, analysis of single-firing BK-sensitive neurons only (n=8 cells, #p<0.05, main effect of BK treatment; two-way mixed ANOVA); right, input-output curves of repetitive-firing neurons (n=7 cells). Data represent mean ± s.e.m.

**Table 3.**
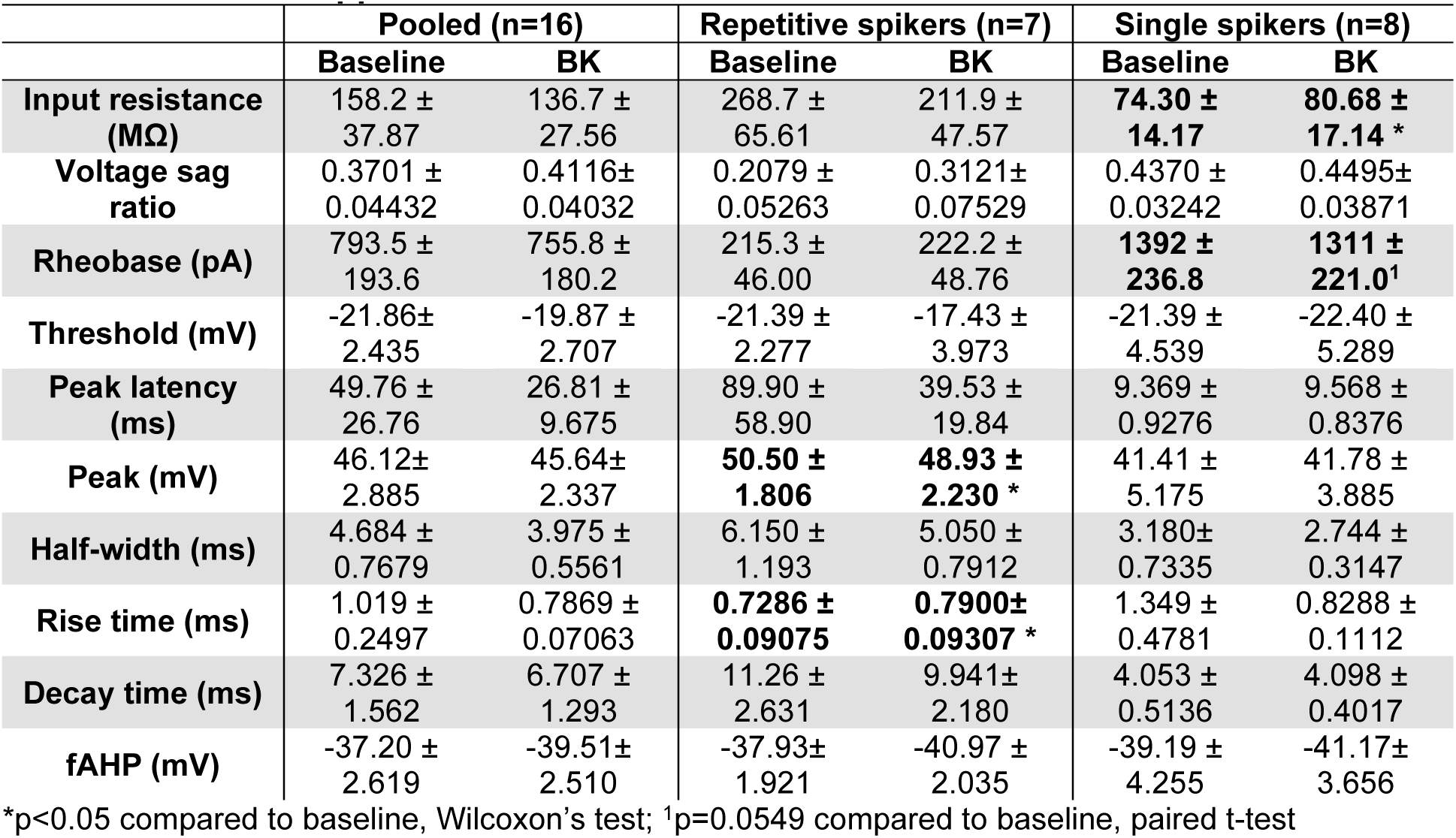
Action potential kinetics and membrane properties of BK-sensitive hDRG before and after acute BK application.

|  | Pooled (n=16) |  | Repetitive spikers (n=7) |  | Single spikers (n=8) |  |
| --- | --- | --- | --- | --- | --- | --- |
|  | Baseline | BK | Baseline | BK | Baseline | BK |
| **Input resistance (MΩ)** | 158.2 ± 37.87 | 136.7 ± 27.56 | 268.7 ± 65.61 | 211.9 ± 47.57 | **74.30 ± 14.17** | **80.68 ± 17.14 \*** |
| **Voltage sag ratio** | 0.3701 ± 0.04432 | 0.4116 ± 0.04032 | 0.2079 ± 0.05263 | 0.3121 ± 0.07529 | 0.4370 ± 0.03242 | 0.4495 ± 0.03871 |
| **Rheobase (pA)** | 793.5 ± 193.6 | 755.8 ± 180.2 | 215.3 ± 46.00 | 222.2 ± 48.76 | **1392 ± 236.8** | **1311 ± 221.0<sup>1</sup>** |
| **Threshold (mV)** | -21.86 ± 2.435 | -19.87 ± 2.707 | -21.39 ± 2.277 | -17.43 ± 3.973 | -21.39 ± 4.539 | -22.40 ± 5.289 |
| **Peak latency (ms)** | 49.76 ± 26.76 | 26.81 ± 9.675 | 89.90 ± 58.90 | 39.53 ± 19.84 | 9.369 ± 0.9276 | 9.568 ± 0.8376 |
| **Peak (mV)** | 46.12 ± 2.885 | 45.64 ± 2.337 | **50.50 ± 1.806** | **48.93 ± 2.230 \*** | 41.41 ± 5.175 | 41.78 ± 3.885 |
| **Half-width (ms)** | 4.684 ± 0.7679 | 3.975 ± 0.5561 | 6.150 ± 1.193 | 5.050 ± 0.7912 | 3.180 ± 0.7335 | 2.744 ± 0.3147 |
| **Rise time (ms)** | 1.019 ± 0.2497 | 0.7869 ± 0.07063 | **0.7286 ± 0.09075** | **0.7900 ± 0.09307 \*** | 1.349 ± 0.4781 | 0.8288 ± 0.1112 |
| **Decay time (ms)** | 7.326 ± 1.562 | 6.707 ± 1.293 | 11.26 ± 2.631 | 9.941 ± 2.180 | 4.053 ± 0.5136 | 4.098 ± 0.4017 |
| **fAHP (mV)** | -37.20 ± 2.619 | -39.51 ± 2.510 | -37.93 ± 1.921 | -40.97 ± 2.035 | -39.19 ± 4.255 | -41.17 ± 3.656 |

\*p<0.05 compared to baseline, Wilcoxon's test; <sup>1</sup>p=0.0549 compared to baseline, paired t-test

Previously, we found that acute BK can lead to changes in action potential kinetics in hDRG neurons [11]. Therefore, we examined physiological phenotype-specific changes in action potential kinetics following acute treatment with BK. Intriguingly, we found that changes in AP kinetics were only seen in repetitively-firing neurons (Fig 4, Table 3), a phenotype associated with a wider AP half-width and slower decay in hDRG [60]. In repetitive spikers (n=8), acute BK led to a slower rise time and a faster decay (Fig 4A-D) and reduced the size of the action potential peak (Fig 4E), consistent with our previous findings [11]. This shift in kinetics was not observed in single-spiking neurons (n=10; Fig 4F-J). Together, this suggests that acute BK may have phenotype-specific effects on AP kinetics.

**Figure 4.**
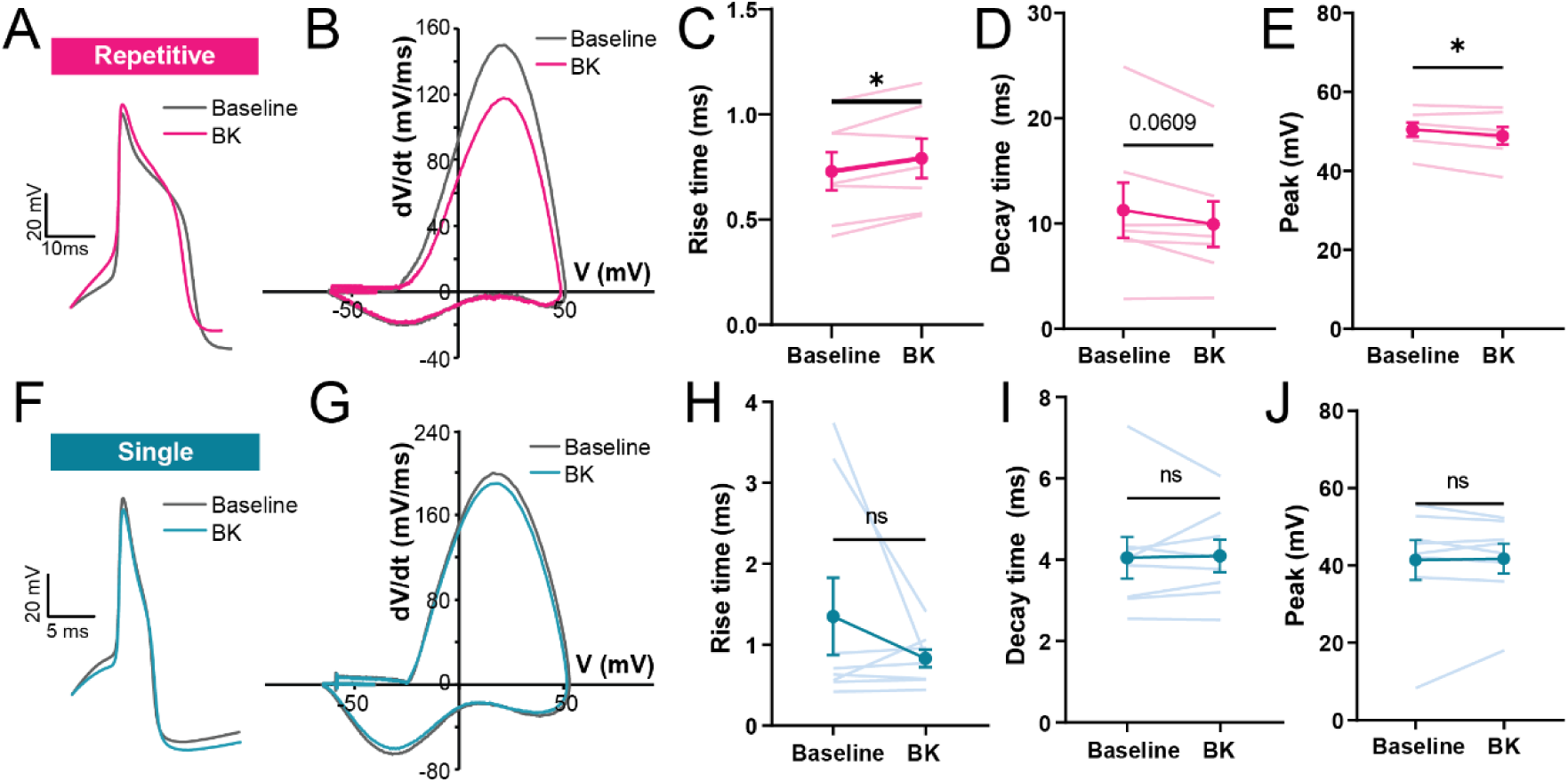
Acute exposure to BK shifts action potential kinetics of repetitive-firing but not single-spiking human DRG neurons. (A) Example voltage trace of an action potential at rheobase of a BK-sensitive, repetitive-firing neuron, before (gray) and after (pink) acute BK (100nM); scale bars, 20mV/10ms. (B) Representative phase-plane plot of a hDRG neuron in (A) at rheobase, before and during application of BK. (C-E) Changes in rise time, decay time, and peak voltage in action potential spikes of repetitive firing neurons, before and during acute BK application; n= 7 cells, *p<0.05, paired t-test. (F) Example voltage trace of an action potential at rheobase of a BK-sensitive, single-spiking neuron, before (gray) and after (blue) acute BK (100nM); scale bars, 20mV/10ms. (G) Representative phase-plane plot of the single-spiking hDRG neuron in (F) at rheobase, before and during application of BK. (H-J) Rise time, decay time, and peak of action potentials from single-spiking neurons before and during acute BK application; n=8 cells. Data represent mean ± s.e.m.

### Prolonged exposure to bradykinin reduces the excitability of repetitive-firing, but not single-spiking, human neurons

Persistent upregulation in BK levels has been implicated in painful inflammatory disorders in humans, such as arthritis [6,21,41] and endometriosis [37]. Therefore, we examined how prolonged (18-24hr) exposure to BK modulates the intrinsic excitability of human DRG neurons. Overnight treatment with BK did not significantly change the proportions of intrinsic firing phenotypes (single vs. repetitive) of neurons recorded (Fig 5A). Among repetitive neurons, we observed a 50% reduction in the relative proportion of neurons that displayed spike frequency adaptation in BK-treated neurons compared to vehicle-treated neurons; however, this difference was not statistically significant (p=0.072, chi-square test).

**Figure 5.**
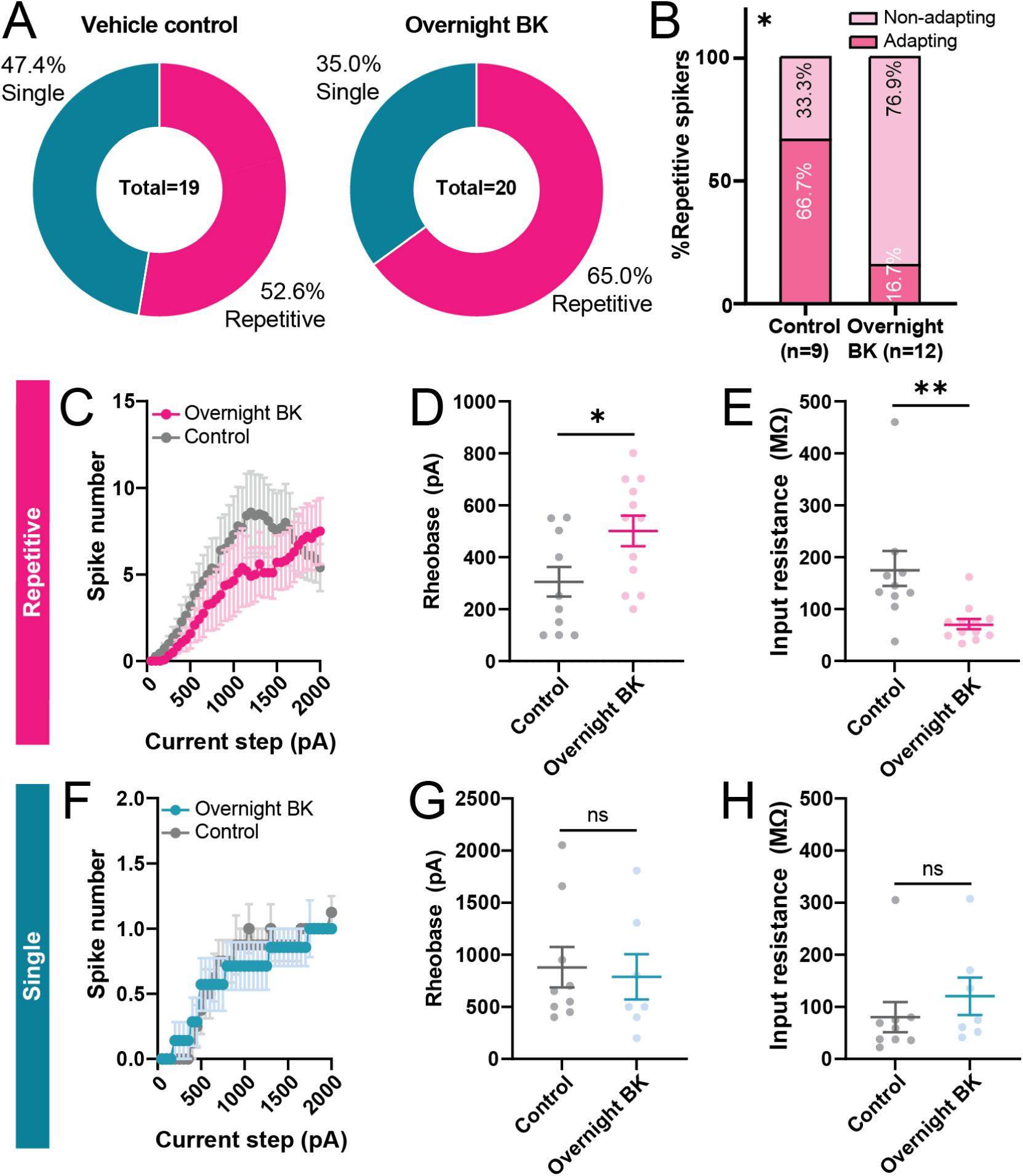
Prolonged exposure to BK leads to reduced excitability in repetitive, but not single, spiking neurons. (A) Distribution of firing patterns of neurons treated overnight with vehicle or BK (100 nM). (B) Relative frequency of repetitive spikers that have adapting and non-adapting firing patterns; *p=0.0195, chi-square test. (C-E) Input-output curve (C), rheobase (D), and input resistance (E) of repetitive-spiking neurons treated with BK or vehicle overnight; n=10-12 cells, *p<0.05, **p<0.01, unpaired t-test. (F-H) Input-output curve (F), rheobase (G), and input resistance (H) of single-spiking neurons treated with BK or vehicle overnight; n=7-9 cells. Data represent mean ± s.e.m.

Among repetitive-spiking neurons, prolonged exposure to BK led to an overall reduction in excitability (Fig 5C-E), including a significant increase in rheobase and reduction in input resistance. We did not observe any difference in action potential kinetics between overnight vehicle- and BK-treated groups (Table 4). Single spikers from vehicle-treated and overnight BK-treated groups did not display significant differences in measures of excitability or spike kinetics (Fig 5F-H, Table 4). Finally, overnight treatment with BK did not significantly reduce the viability of cells compared to vehicle (74.68% vs. 77.06%, Vehicle vs. BK, p=0.86, unpaired t-test) indicating that observed physiological changes were not likely due to selective cell death of subpopulations of neurons following BK treatment. Together, these results suggest that prolonged BK treatment modulates the excitability of human DRG neurons in a physiological phenotype-dependent manner.

**Table 4.**
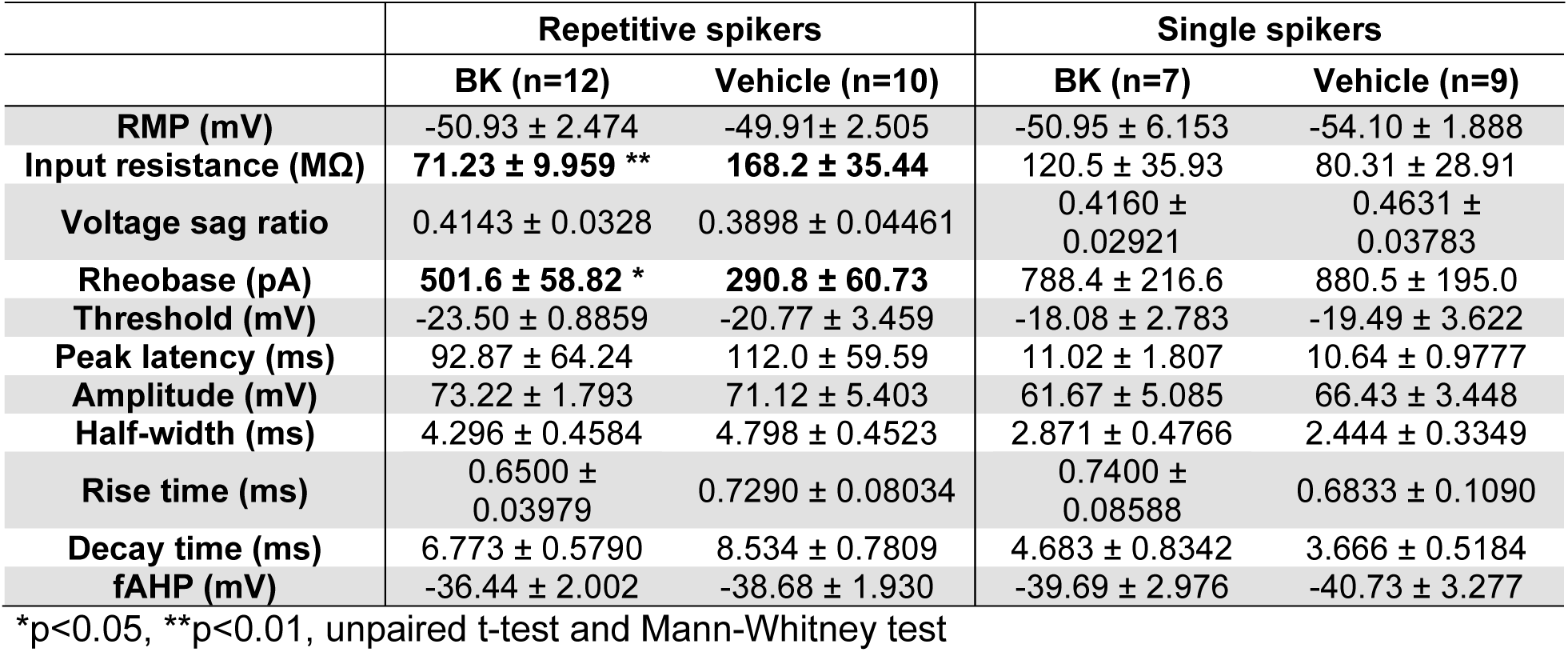
AP kinetics and membrane properties of overnight vehicle- and BK-treated cells.

|  | Repetitive spikers |  | Single spikers |  |
| --- | --- | --- | --- | --- |
|  | BK (n=12) | Vehicle (n=10) | BK (n=7) | Vehicle (n=9) |
| **RMP (mV)** | -50.93 ± 2.474 | -49.91 ± 2.505 | -50.95 ± 6.153 | -54.10 ± 1.888 |
| **Input resistance (MΩ)** | **71.23 ± 9.959 \*\*** | **168.2 ± 35.44** | 120.5 ± 35.93 | 80.31 ± 28.91 |
| **Voltage sag ratio** | 0.4143 ± 0.0328 | 0.3898 ± 0.04461 | 0.4160 ± 0.02921 | 0.4631 ± 0.03783 |
| **Rheobase (pA)** | **501.6 ± 58.82 \*** | **290.8 ± 60.73** | 788.4 ± 216.6 | 880.5 ± 195.0 |
| **Threshold (mV)** | -23.50 ± 0.8859 | -20.77 ± 3.459 | -18.08 ± 2.783 | -19.49 ± 3.622 |
| **Peak latency (ms)** | 92.87 ± 64.24 | 112.0 ± 59.59 | 11.02 ± 1.807 | 10.64 ± 0.9777 |
| **Amplitude (mV)** | 73.22 ± 1.793 | 71.12 ± 5.403 | 61.67 ± 5.085 | 66.43 ± 3.448 |
| **Half-width (ms)** | 4.296 ± 0.4584 | 4.798 ± 0.4523 | 2.871 ± 0.4766 | 2.444 ± 0.3349 |
| **Rise time (ms)** | 0.6500 ± 0.03979 | 0.7290 ± 0.08034 | 0.7400 ± 0.08588 | 0.6833 ± 0.1090 |
| **Decay time (ms)** | 6.773 ± 0.5790 | 8.534 ± 0.7809 | 4.683 ± 0.8342 | 3.666 ± 0.5184 |
| **fAHP (mV)** | -36.44 ± 2.002 | -38.68 ± 1.930 | -39.69 ± 2.976 | -40.73 ± 3.277 |

\*p<0.05, \*\*p<0.01, unpaired t-test and Mann-Whitney test

### Documented pain history and BK receptor expression or BK-mediated sensitization

Lasting upregulation in B1 and B2 receptor expression and increase in the production of BK have been associated with various rodent models of pain, including myalgia, neuropathic pain, and inflammatory pain [2,15,24,26,38,45,59]. Therefore, we examined whether history of pain was associated with differences in BK receptor expression in the hDRG. First, we analyzed a previously published bulk RNA-seq dataset of hDRG obtained from cancer patients with or without radicular/neuropathic pain [42]. Whole-tissue level of *BDKRB1* was ∼1.5 fold higher in hDRG from patients with radicular/neuropathic pain, though this difference was not significant (p=0.3539, Mann-Whitney test) (Fig 6A). *BDKRB2* level in whole-DRG samples was comparable between the two groups (p=0.6026, Mann-Whitney test) (Fig 6B).

**Figure 6.**
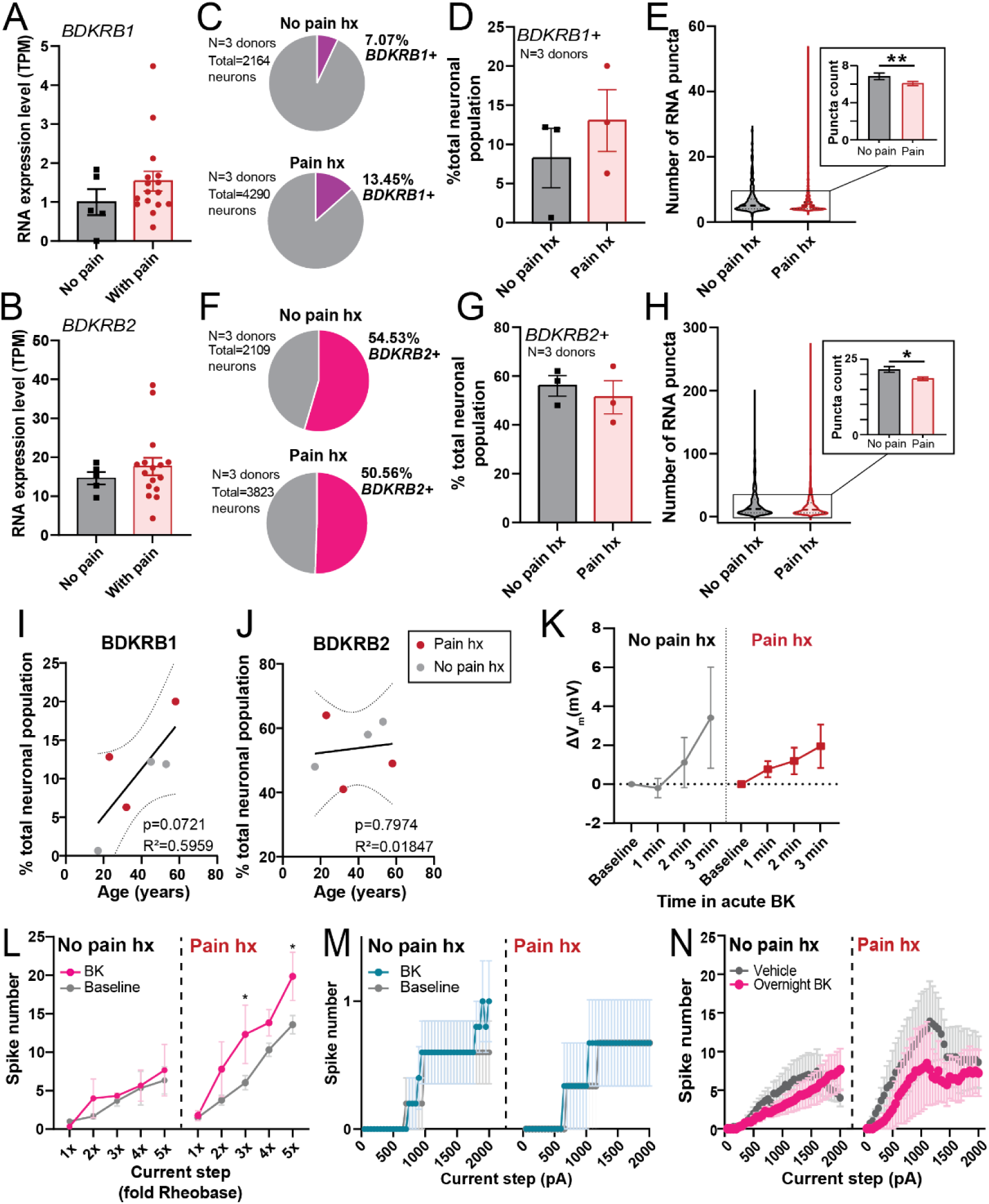
BK receptor expression and BK-mediated sensitization in donors with documented pain history. Whole-tissue *BDKRB1* (A) and *BDKRB2* (B) expression level, in TPM, in hDRG from cancer patients with or without neuropathic pain (data re-analyzed from North et al., 2019 [43]). (C) Pie-chart showing relative proportion of *BDKRB1*-positive hDRG neurons in a pooled sample of hDRG neurons (n= 2164-4290 from 3 donors per group; 1207-1632 neurons per each donor in the “Pain hx” group, 300-950 neurons per each donor in the “No pain hx” group). (D) Percentage of *BDKRB1-*expressing DRG neurons per individual donor (N=3 donors, 2-3 sections and 323-1632 cells per donor). (E) Distribution of *BDKRB1* RNA expression within each *BDKRB1*-positive cell. Dotted line on violin plot indicates median number of *BDKRB1* puncta per neuron. Inset shows a bar graph of the mean puncta count per cell for *BDKRB1* (n=152-429 *BDKRB1+* neurons from 3 donors for each condition; **p<0.01, Mann-Whitney test). (F) Pie-chart displaying relative proportion of *BDKRB2*-positive hDRG in a pooled sample of hDRG neurons (n=2109-3823 neurons pooled from 3 donors per group; 963-1890 neurons per donor in the “Pain hx” group, 340-1044 neurons per donor in the “No Pain hx” group). (G) Percentage of *BDKRB2*-expressing DRG neurons per individual donor (N=3 donors, 338-1890 cells per donor). (I) Distribution of *BDKRB2* RNA expression level within each *BDKRB2*-positive cell. Dotted line on violin plot indicated median number of *BDKRB2* puncta per neuron. Inset shows a bar graph of the mean puncta count per cell for *BDKRB2* (n=655-1542 *BDKRB2+* neurons from 3 donors for each condition; *p<0.05, Mann-Whitney test). (I) Percentage of *BDKRB1-*expressing hDRG neurons per donor plotted against donor’s age. Line shows line of best fit predicted by simple linear regression analysis (Y = 0.3050*X - 0.9456), ± standard error of the slope (dotted line). (J) Percentage of *BDKRB2-*expressing hDRG neurons per donor plotted against donor’s age. Solid line shows line of best fit predicted by simple linear regression analysis (Y = 0.07413*X + 50.85), ± standard error of the slope (dotted line). (K) Change in resting membrane potential over time during acute BK application in hDRG neurons from donors with (right; n=5) and without (left; n=15) history of pain. (L) Input-output curves of BK-sensitive repetitive-spiking hDRG neurons from donors with (right; n=4) and without (left; n=3) history of pain, before and during acute BK treatment. *p<0.05, two-way repeated measures ANOVA with Šídák’s post hoc test. (M) Input-output curves of BK-sensitive single-spiking hDRG neurons from donors with (right; n=3) and without (left; n=5) history of pain, before and during acute BK treatment. (N) Effect of overnight BK or vehicle on input-output curves of hDRG neurons from donors with and without documented pain history (n=3-8 per group). Data represent mean ± s.e.m.

As whole-DRG RNA-seq does not distinguish between neurons and non-neuronal cells in the tissue, we next assessed whether there are neuron-specific differences in the expression of BK receptors in the hDRG from donors with and without history of pain. We identified three donors with documented pain history in their medical history (Table 2) and assessed expression of B1 and B2 receptors in their DRGs using RNAScope *in situ* hybridization. We observed that the proportion of hDRG neurons that expressed *BDKRB1* was 1.9-fold higher in the “pain history” group compared to the “no pain history” group (Fig 6C, D), though this difference was not statistically significant (p=0.4313, unpaired t-test). Within *BDKRB1*-expressing cells, pain history was associated with a significant but small decrease in the average *BDKRB1* RNA expression level per cell (p<0.01, η^2^=0.018, d=0.228; Fig. 6E). Proportion of BDKRB2-expressing hDRG neurons was comparable between the two groups (p=0.5875; Fig 6F, G). While average expression level of *BDKRB2* within each *BDKRB2*+ neuron was significantly lower in the pain history group, the effect size of this difference was trivial (p<0.05, η^2^=0.002, d=0.098; Fig 6H). Interestingly, we observed a positive linear relationship approaching significance (p=0.0721, R^2^=0.5959) between age and proportion of *BDKRB1*-expressing hDRG neurons (Fig 6I), but not in *BDKRB2-*expressing neurons (Fig 6J).

Next, we tested whether pain history was associated with any differences in the overall modulatory effects of BK on neuronal physiology. To that end, we re-analyzed the physiology experiments (Fig 3-5) based on donors’ pain history (Table 1). Acute BK led to depolarization in hDRG neurons from both donors with and without pain history (Fig 6K). The hyperexcitability phenotype of acute BK appeared more robust in repetitively-spiking hDRG neurons obtained from donors with pain history (Fig 6L); effect of acute BK on input-output curves of single-spiking neurons was comparable across both pain groups (Fig 6M). We were unable to assess whether pain history was associated with difference in the relative magnitude of acute BK-induced hyperexcitability in repetitive spikers due to the small sample size per group resulting from segmentation by both pain history and spiking phenotype (n=3-5 per spiking phenotype per pain group), Finally, overnight BK treatment led to reduced spike firing and excitability in repetitive neurons from both donors with and without pain history (n=3-8; Fig 6N, Table 5). Together, this suggests that modulatory effect of acute and prolonged BK on neuronal excitability is broadly consistent in sensory neurons obtained from donors with and without pain history.

**Table 5.** Rheobase and input resistance of overnight vehicle- and BK-treated repetitive spiking neurons, segmented by donors’ pain history.

|  | No pain history |  | Pain history |  |
| --- | --- | --- | --- | --- |
|  | BK (n=8) | Vehicle (n=8) | BK (n=4) | Vehicle (n=3) |
| **Rheobase (pA)** | 483.2 ± 68.33 | 369.7 ± 63.60 | 538.5 ± 124.9<sup>1</sup> | 133.5 ± 33.45 |
| **Input resistance (MΩ)** | 76.82 ± 14.09* | 190.0 ± 46.26 | -60.06 ± 10.10** | 147.5 ± 9.114 |

\*p<0.05, Mann-Whitney test compared to Vehicle; \*\*p<0.01, unpaired t-test compared to Vehicle; <sup>1</sup>p=0.0571, Mann-Whitney test compared to Vehicle.

## Discussion

In the present study, we show that human nociceptors express functional BK receptors, and that BK can bidirectionally modulate the membrane excitability of hDRG neurons depending on the duration of exposure and the physiological subtype of the neuron. Our results also suggest that pain history may be associated with higher B1 receptor expression in hDRG and more robust neuronal response to acute BK. Together, these experiments support the notion that BK signaling is a potential analgesic target for inflammatory pain in humans.

In rodents, BK’s hyperalgesic effect on naïve animals is mediated by the B2 receptors expressed on peptidergic nociceptors [9,12,34,36,54]. Consistent with these findings in rodents, we found that B2 receptors are particularly enriched among putative *SCN10a+* nociceptors in the human DRG; we did not distinguish between peptidergic and non-peptidergic populations of human nociceptors in our study, as human DRG, unlike rodent DRG, has a high overlap between these markers [51].

The present findings support and extend our previous reports that hDRG neurons can be activated and/or sensitized by acute BK [11]. The hyperexcitability-inducing effects of acute BK may partly account for the increases in pain rating seen in healthy human volunteers that receive intradermal BK injection, and in other inflammatory disorders like arthritis that are associated with elevated plasma and synovial kinin levels [6,21,33]. One possible mechanism underlying acute BK-induced hyperexcitability is G_q_-mediated inhibition of M-type K^+^ currents and activation of the Ca^2+^-gated chloride channel, ANO1, both of which contribute to BK-induced membrane depolarization and spike firing in rat DRG [28,58]. The fact that multiple subpopulations of hDRG neurons express the genes for both Kv7 channels and ANO1 [55] further bolsters this idea, though additional functional studies are needed. Together, these results indicate that peripheral mechanism underlying acute BK-induced pain may be conserved from rodents to humans.

Overnight incubation with BK and other inflammatory reagents has been previously used to mimic chronic inflammatory disorders like arthritis *in vitro* [22,54]. Surprisingly, prolonged BK led to a reduction in the excitability in a subset of hDRG neurons. The reduction in excitability in neurons with a repetitive spiking phenotype was also accompanied by a partial increase in the incidence of repetitive spikers without spike frequency adaptation, a cluster associated with smaller number of action potentials discharged at lower current steps [60] Similar shift towards a hypoexcitable firing phenotype has been seen previously in human DRG neurons following prolonged depolarization[47]; and in DRG from rats with spinal cord injury that were exposed *in vitro* to high concentrations of the cytokine macrophage migration inhibitory factor [3]. In conjunction with these studies, our findings provide additional evidence that human sensory neurons can engage in a homeostatic shift towards a more hypoexcitable state in the context of prolonged inflammation.

A large number of preclinical studies has shown that pain-inducing injury and inflammation can upregulate B2 receptor levels and induce B1 receptor synthesis in the rodent DRG [12,13,24,26,30,38,44,57,59]. In this study, we attempted to determine whether donor’s prior history of pain similarly correlates with differential levels of BK receptor expression. Usage of tissue from human organ donors presents several limitations, including (a) variation in pain phenotype; (b) inability to test which neurons within the DRG innervate the site of pain; (c) large variability in demographic and medical histories within the donor pool; and (d) limited availability of samples. Despite these sources of variability, our analyses suggest that donor’s pain history may be associated with greater number of B1 receptor-expressing neurons in the hDRG; this is also supported by our functional data that suggests that in repetitive-spiking neurons from donors with pain history, acute BK has a more robust effect on neuronal excitability. If pain-inducing injury or inflammatory insult leads to de novo synthesis of B1 receptors at low levels in hDRG neurons, then this would account for the seemingly opposite trends of greater percentage of *BDKRB1*+ neurons but lower average within-cell expression level of *BDKRB1* in hDRG from pain donors. These trends may also explain why the relative increase in B1 receptor expression in the pain history group is much smaller in the whole-DRG transcriptomics dataset from cancer patients[42], as whole tissue transcriptomics cannot distinguish between differing expression levels in multiple neurons. Additionally, the difference in the pain pathology of the donors in the two datasets (radicular/neuropathic/cancer vs. predominantly musculoskeletal), as well as the inclusion of non-neuronal cells in the whole transcriptomics analysis, may additionally contribute to the differences in the results of the two analyses.

We were surprised to see a positive linear trend between age and proportion of B1 receptor-expressing neurons in the hDRG, which suggests that age alone may also be an important factor associated with BK receptor expression. Age-related increases in B1 and/or B2 receptor levels have been previously reported in rat and human cardiac tissue [23,27]. A larger dataset and sample pool would allow us to more effectively conclude whether pain history, pathology, and age are significant predictors of B1 receptor expression among hDRG neurons at a population level.

Previous studies have shown that satellite glial cells in rodent DRG and trigeminal ganglia express functional BK receptors and appear to play a role in BK-induced inward current in rat sensory neurons [7,20]. Our results demonstrate that B2 receptors are also expressed on human satellite glial cells. This finding raises the possibility that satellite glial cells may be involved in BK-mediated neuronal hyperexcitability in hDRG. Recent studies have shown that satellite glial cells in mouse and rat DRG can release inflammatory interleukins, PGE2, and TNF-α, which can, in turn, induce hyperexcitability in DRG neurons [31,53]. Whether activation of B2 receptors on satellite glial cells can trigger similar inflammatory glia-neuronal signaling is an interesting avenue for further research.

Although many studies in rodents have previously shown that BK antagonists can be analgesic across multiple different models of pain [10,13,16,18,38,40,59], there have been few published clinical studies assessing the efficacy of BK receptor antagonists in human pain patients. Icatibant (HOE-140) is a B2 receptor antagonist that has been FDA-approved for the treatment of hereditary angioedema (HAE), a rare and painful inflammatory disorder. While BK signaling has been implicated in multiple other painful disorders (notably, arthritis) in human studies [6,21,37], the progress towards repurposing this orphan drug for other inflammatory or generalized pain conditions has been slow. For example, according to the NIH’s database on ClinicalTrails.gov, only one clinical study has been conducted since 2000 to study icatibant’s efficacy against pain in non-HAE patients. A study published in 2008 [52] found that arthritis patients who received intra-articular injections of icatibant experienced a significant improvement in pain at rest and during activity compared to the placebo group. Despite the small sample size, results of this study provide promising evidence of analgesic efficacy of BK receptor antagonists in a wider range of painful inflammatory disorders [52].

Altogether, our findings suggest the mechanism underlying BK-mediated pain and sensitization are generally conserved from rodents to human. The consistency in BK receptor expression patterns and its acute effects across species makes BK signaling an ideal target for further study and drug development that could be probed using both preclinical (e.g. rodent) and translational (e.g. primary human neuronal culture) approaches. In the context of previous studies in rodents and the clinical work outlined above, our study further supports the notion that BK signaling in the peripheral nervous system is a viable target for pain relief in multiple disorders in humans.

## Methods

### Human DRG extraction

hDRG extraction and culture was performed as previously described [56]. Briefly, T11-L5 DRGs were extracted from postmortem organ donors in collaboration with Mid-America Transplant. Donor’s medical history, including history of chronic pain, was determined based on Mid-America Transplant’s interview with the donor’s family (Table 1, 2). Extraction was performed within 2 hours of aortic cross-clamp and hDRG samples were placed in ice-cold N-methyl-D-glucamine (NMDG)-based solution (93mM NMDG, 2.5mM KCl, 1.25mM NaH_2_PO_4_, 30mM NaHCO_3_, 20mM HEPES, 25mM glucose, 5mM ascorbic acid, 2mM thiourea, 3mM Na^+^ pyruvate, 10mM MgSO_4_, 0.5mM CaCl_2_, 12mM N-acetylcysteine; adjusted to pH7.3 using NMDG or HCl, and 300-310mOsm using H_2_O or sucrose) for transport. Extracted hDRG were cleaned and cultured in lab immediately following extraction.

### Human DRG culture

Several thoracolumbar hDRGs were pooled for each culture. Following previously published protocol for hDRG culture[56], hDRGs were first enzymatically dissociated with papain (Worthington) and collagenase type 2 (Sigma), then mechanical dissociated by trituration. Dissociated DRGs were filtered using a 100 μm cell strainer (Fisher) and cultured on collagen-coated glass coverslips in DRG media consisting of Neurobasal A medium (Gibco), 5% fetal bovine serum (Gibco), 1% penicillin/streptomycin (Corning), Glutamax (Life Technologies), and B27 (Gibco). hDRG were used for experiments at days in vitro (DIV) 4-10.

### RNAScope *in situ* hybridization (ISH)

Lumbar hDRG not used for culture were fixed in 4% paraformaldehyde overnight, transferred to solution containing 30% sucrose for 10 days or until the tissue sank to the bottom of the container, embedded in OCT in a cryomold, flash-frozen, and stored at -80°C until use. 12-μm sections were prepared from fixed-frozen hDRG tissue, mounted on Superfrost Plus slides, and stored at -20°C or -80°C with desiccant until used.

Slides containing hDRG sections were dehydrated using 50, 70, and 100% ethanol, and treated with Protease IV and hydrogen peroxide (ACDBio). The following probes from ACDBio were used: BDKRB1 (#424321), BDKRB2 (#424331-C2), SCN10A (#406291-C3). RNAScope was performed following manufacturer’s protocol. At the end of RNAScope, TrueBlack® Autofluorescence Quencher (Biotium) was applied to sections as directed by manufacturer protocol. Slides were subsequently washed 3x with PBS. DAPI was applied to the slides immediately before or after the application of TrueBlack®. Slides were mounted with coverslips using ProLong Gold Antifade Mountant (Invitrogen) and stored at -20°C. Slides were imaged within 3 days.

### Immunohistochemistry (IHC)

For experiments involving dual ISH-IHC, IHC was performed after RNAScope. After the final HRP signal was developed, slides were washed 2 times in PBS-T (PBS with 0.1% Tween-20), then incubated in blocking buffer (10% donkey serum in PBS-T) for 1 hour at room temperature in the dark. Slides were then incubated in primary antibody (rabbit anti-FABP7; Invitrogen) diluted 1:500 in blocking buffer overnight at 4°. The next day, slides were washed 3 times in PBS-T and incubated in secondary antibody (goat anti-rabbit 647; Invitrogen) diluted 1:1000 in PBS for 2 hours at room temperature. Slides were washed 3 times in PBS before application of DAPI, treatment with TrueBlack, and mounting as described above.

### Imaging

Following RNAscope, slides were imaged using Leica DM6b system at 20x magnification. The entire human DRG section was imaged and merged using Leica Application Suite X (LAS-X, v.3.7, Leica Microsystems). Acquisition setting were kept consistent throughout imaging across different experimental days. Acquisition settings for each fluorescence channel were as follows: DAPI, 50ms exposure, 2.1 gain, FIM 55%; L5, 300ms exposure, 2.5 gain, FIM 55%; TXR, 500 ms exposure, 3.0 gain, FIM 55%; and Y5 500 ms exposure, 2.0 gain, FIM 55%.

### Image analysis

For analysis, 2-3 hDRG sections per donor were chosen at random, though preference was given to sections that did not have extensive artifacts from sectioning and mounting, such as tear in section or bubbles. DRG cell body profiles were visually identified using autofluorescence and DAPI fluorescence from the satellite glial cells surrounding neuronal cell bodies using methods described previously [51]. Identified cell bodies were manually segmented using the freehand ROI tool in ImageJ (NIH). The number of cells per section varied between sections, ranging from 100 to 900. For each cell, its area was measured using ImageJ and equivalent diameter was calculated using the formula, 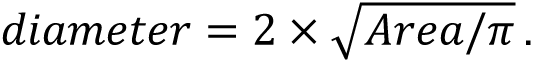 Following segmentation, cells were manually classified as “positive” or “negative” for each gene of interest based on the presence of RNA puncta in the soma. Fluorescent signal that appeared across all 3 channels were considered as background aurofluorescence and not included in analysis. For each experiment, any adjustment to the brightness and contrast were kept consistent across all sections and donors used for that experiment. Proportion of neurons expressing the gene of interest was calculated by dividing the sum of all “positive” neurons in all the sections for each donor by the sum of all neurons in all sections for that specific donor.

### Patch clamp electrophysiology

At DIV 4-10, coverslips containing cultured primary hDRG neurons were transferred to a recording chamber. Experiments were conducted in external solution containing (in mM): 145 NaCl, 3 KCl, 2 CaCl_2_, 1 MgCl_2_, 7 glucose, 10 HEPES, adjusted to pH 7.3 with NaOH. Whole-cell patch-clamp recordings were made using fire-polished borosilicate glass pipettes with 2-3.5 MΩ resistance. The pipettes were filled with K-gluconate based internal solution consisting of the following (in mM): 120 K-gluconate, 5 NaCl, 3 MgCl_2_, 0.1 CaCl_2_, 10 HEPES, 1.1 EGTA, 4 Na_2_ATP, 0.4 Na_2_GTP, 15 Na_2_Phosphocreatine; adjusted to pH 7.3 with KOH and HCl, and 290 mOsm with sucrose. All experiments were conducted at room temperature. External solution was perfused continuously using gravity-fed bath perfusion at a flow rate of 1-2mL/min.

Recordings were made using a MultiClamp 800B amplifier and a Digidata 1550B digitizer (Molecular Devices, CA). Data were sampled at 10 kHz. Electrophysiology data were compiled using ClampFit (v.11.1, Molecular Devices). Series resistance was kept below 15 MΩ in all current clamp recordings.

After a stable whole-cell configuration was achieved, membrane excitability was assessed in current-clamp mode. Cells whose resting membrane potentials were below -75mV or above -35 mV were considered unhealthy and excluded from analysis. A small number of neurons displayed spontaneous activity at rest that prevented us from measuring their resting potential. These spontaneously active neurons were included for analysis if they required less than 500pA of negative holding current to be held at -60mV. Active and passive membrane properties were analyzed using Easy Electrophysiology (Easy Electrophysiology Ltd., UK). Resting membrane potential was continuously monitored throughout the experiment. Input-output relationship was determined by counting the number of action potentials generated by 1-second-long current steps from 50pA to 2nA (Δ50pA). Rheobase was defined as the minimum amount of current step required to evoke an action potential.

Action potential kinetics were analyzed from the first action potential evoked at rheobase. Threshold was defined as membrane voltage when dV/dt = 10. AP rise-time and decay-time were defined as time from 10-90% of rising phase or of the falling phase, respectively. Half-width was measured as the time between 50% of the rise and decay phase of the action potential. AP amplitude was measured from the AP threshold to peak. Passive membrane properties, including input resistance and voltage sag, were measured using 1-second-long hyperpolarizing current steps from -250 to -50pA (Δ50pA) from a holding potential of -60mV. For experiments assessing effects of acute BK, the above measures were assessed both before and after application of BK.

### Classification of firing phenotype

Cells were categorized as single- or repetitively-spiking. Cells that fired no more than one action potential in response to depolarizing current steps from 50pA to 2nA (Δ50pA) and at 2-3x rheobase were categorized as “single-spiking,” similar to the “rapidly adapting” clusters described in rat DRG [3,43]. Cells which fired more than one action potential across multiple stepwise current injections were categorized as “repetitive-spiking” neurons, similar to the “non-adapting” clusters identified in rat DRG [3,43]. Repetitive-spiking neurons were further categorized into adapting and non-adapting clusters based on the presence of spike frequency adaptation. A cell was classified as “repetitive adapting” if the cell displayed an inverse U-shaped input-output curve of spike number vs. current injected; while cells displaying a linearly increasing input-output curve with no spike frequency adaptation were classified as “repetitive non-adapting.”

### Viability assay

Cultured human DRG neurons were treated with either 100 nM BK, 200 mM KCl, or vehicle (0.1mM acetic acid) overnight (18-24 hrs). Following overnight treatment, plated DRGs were harvested by scraping the coverslip with p1000 pipette tips. Cell suspension was collected and spun down at 1000 rpm for 3min. Supernatant was discarded, and the pellet was re-suspended in 20-50 μL of fresh HBSS + HEPES buffer. Calcein-AM (Invitrogen) stock was prepared using DMSO according to manufacturer’s instructions. A 10 μM working solution of Calcein-AM was prepared in HBSS+H. The cell suspension (10 μL) was diluted with equal volume of Calcein-AM and loaded onto a hemocytometer. The cells were imaged using Leica DM6b system with LAS-X (v.3.7, Leica Microsystems) at 20x. Presence of green fluorescence was utilized to distinguish between live (fluorescent) and dead (non-fluorescent) cells. Viability ratio was calculated by dividing the number of live cells over the total number of cells (sum of live and dead cells).

### Pain history assessment

Donors’ pain history was assessed by reviewing the social and medical history of the donor as documented on the United Network for Organ Sharing (UNOS) records. Donors whose UNOS records contained key words such as “constant pain,” “frequent pain,” “joint pain,” and/or diseases associated with pain (e.g. scleroderma, arthritis) were designated as having a prior history of chronic or persistent pain (Table 1, 2).

### Statistics

Statistical analyses were performed using GraphPad Prism 9 (GraphPad Software, LLC). For each dataset analyzed, normality of residuals was tested using the Shapiro-Wilk test to inform the use of parametric or non-parametric tests. To assess statistical differences between two groups, t-test (paired or unpaired; as indicated in text), Mann-Whitney tests, or Wilcoxon’s tests were performed. To assess statistical differences across multiple groups and/or time-points, one- or two-way repeated measures ANOVAs with Geisser-Greenhouse correction and Sidak’s post hoc test, Friedman’s test with Geisser-Greenhouse correction and Dunn’s posthoc test, or restricted maximum likelihood (REML) mixed effects analysis with Geisser-Greenhouse correction and Sidak’s posthoc tests were used. For select tests with significant results as defined by p<0.05, effect size was calculated[25]. All data are represented as mean ± SEM. Detailed statistics are reported in Supplementary Tables S1-31.

## Supporting information

Supplementary statistics tables S1-31

## Acknowledgements

This work was supported by the American Heart Association through Predoctoral Fellowship #828671 (J.Y.) and by the National Institutes of Health through the NIH HEAL Initiative under award number U19NS130607 (R.G.). We thank the donors, their families, and Mid-America Transplant, without whom this research would not have been possible; J. Lemen for his support during surgical extractions; and the past and present members of the Gereau lab for their helpful comments and critiques.

